# Biosynthesis, Characterization and Anthelmintic Activity of Silver Nanoparticles of *Clerodendrum infortunatum* Isolate

**DOI:** 10.1101/2023.03.20.533573

**Authors:** Rima Majumdar, Pradip Kumar Kar

**Affiliations:** Parasitology Laboratory, Department of Zoology, Cooch Behar Panchanan Barma University, Cooch Behar, West Bengal, India

## Abstract

Over the past few decades, the green synthesis of nanoparticles has gained importance for their therapeutic efficacy and eco-friendly nature. Integrating green chemistry principles into multidisciplinary nanoscience research has paved the way for developing environmentally benign and sustainable methods for synthesizing gold and silver nanoparticles. In the present study, the flowers obtained from *Clerodendrum infortunatum* (L.), belonging to the family Verbenaceae, have been used for biosynthesizing silver nanoparticles (AgNPs) to evaluate the anthelmintic potential. UV-Vis spectroscopy, XRD, FTIR and TEM analyses were performed to ascertain the formation of AgNPs. Clerodendrum-derived AgNP (CLE-AgNP) has significantly affected the normal physiological functions of the poultry parasite *Raillietina* spp, a menace to the livestock industry. Our study manifests that CLE-AgNPs cause considerable distortion of the surface tegument of this cestode parasite leading to changes in the host-parasite interface. The histochemical localization studies of the tegument-associated enzymes viz. AcPase, AlkPase, ATPase and 5’-Nu, exposed to the drug, showed a substantial activity decline, thus establishing the anthelmintic potential of the CLE-AgNPs.

## Introduction

Current trends in animal welfare boost the adoption of organic, pen-free range, and backyard husbandry practices. Local chicken managed under the backyard poultry production sector is critical in providing income for small societies. However, the growth of this sector is greatly hampered by the re-emergence of a diverse array of poultry helminths. Several studies suggest chickens and turkeys serve as hosts to a wide range of helminths, causing a huge economic loss in tropical countries like India. The cestode parasite *Raillietina* spp. is highly prevalent in common domestic fowl *Gallus gallus domesticus*, causing enteritis and weight loss in young chickens^1,2^. Common parasite eradication strategies include using the benzimidazoles (BMZ) class of compounds, such as flubendazole, fenbendazole, and albendazole, whose unregulated usage can lead to antihelmintic resistance^3^. Although ethnoveterinary medicine is a well-established practice, evidence on the pharmacology of plant anthelmintics for use in chickens is limited. The current investigation has been conducted to support the therapeutic use of *Clerodendrum infortunatum* (CLE) for the biosynthesis of AgNP as an anthelmintic against the bird cestode *Raillietina* spp.

Bioinspired technology for nanoparticle (NP) synthesis has become a major branch within nanoscience and nanotechnology. So far, numerous metal NPs and metal oxides have been synthesized using plant extracts, microbes, etc^4,5^. Due to their wide availability, renewability, and environmental friendliness, in addition to their immense applications in the synthesis of NPs, plant biomass is mostly targeted as a catalyst for chemical synthesis and biodiesel production^6-8^. Current research suggests that silver nanoparticles (AgNPs) can be used in various medical applications, including antibacterial, antifungal, anti-diabetic, anti-inflammatory, and cancer treatment, as well as diagnosis^9-14^. Silver products have long been known to have strong inhibitory and bactericidal effects and a broad spectrum of antimicrobial activities, which have been used for centuries to prevent and treat various diseases, most notably infections^15^. The synthesis of silver nanoparticles by physical and chemical routes poses serious problems like high capital investment, usage of hazardous chemicals, high temperature and pressure, and toxic solvents^16-18^. Compared to microorganisms, applying plant extracts to synthesize AgNP is more advantageous in resource availability, security, reaction rate and convenience, and feasibility of large-scale synthesis^19-22^. The phytochemicals present in plant extracts have been reported to cause the reduction of metal ions to nanoparticles and eventually obliterate the use of toxic chemicals, high pressure, temperature, energy and maintenance of microbial cultures^23-27^. Various plant materials, such as leaf extracts, fruit, bark, fruit peels, root and callus, have been explored to synthesize NPs in different sizes and shapes^28^. Tripathi *et al*. evaluated the bactericidal activity of silver nanoballs at a concentration of 40 μg/mL against *Escherichia coli, Salmonella typhimurium, Bacillus subtilis*, and *Pseudomonas aeruginosa* by measuring colony-forming units (CFU)^29^. In a previous study, Kar *et al*. investigated the *in vitro* anthelmintic activity of the nanogold particles synthesized by mycelia-free culture filtrate of the fungus *Nigrospora oryzae* treated with gold chloride on worm parasites using a cestode (tapeworm) model^30^. The study revealed alterations in the enzyme activity and effect on the normal physiological functioning of the parasite after treatment with gold nanoparticles.

In this research work, we focus on the bio-augmented synthesis of AgNPs using an aqueous extract of medicinally important CLE. Clerodendrum is a genus of flowering plants in the Lamiaceae (Verbenaceae) family that is very common throughout the plains of India, found widely in West Bengal. Although above 400 species of the genus, Clerodendrum are distributed worldwide, only a few have been investigated and studied so far^31^. Plants belonging to the genus Clerodendrum are well known for their pesticidal properties, and various Clerodendrum species such as *C. indicum, C. phlomidis, C. serratum var. amplexifolium, C. trichotomum, C. chinense, C. petasites* have been historically used as folk and traditional medicine to treat diseases, such as cold, hyperpyrexia, asthma, furunculosis, hypertension, rheumatism, dysentery, mammitis, toothache, anorexia, leucoderma, leprosy, arthrophlogosis, and other inflammatory diseases in numerous locations around the globe such as India, China, Korea, Japan, Thailand, and Africa^32-34^. The plant contains triterpenes, steroids and flavonoids, and tribes use various parts of the plant in colic, scorpion sting and snake bite, smallpox, tumors and certain skin diseases^35-38^. The current work intends to explore the *in vitro* anthelmintic activity of AgNPs generated from the aqueous floral extract of CLE on the cestode *Raillietina* spp. The study will evaluate the efficacy of the AgNPs as a potential anthelmintic treatment and contribute to understanding the mechanisms underlying their anthelmintic activity. The results of this study may provide new insights into the development of sustainable and eco-friendly treatments for helminth infections and help to address the problem of drug resistance in current treatments.

## Results

### Characterization studies

#### UV-Visible Spectrophotometric Analysis

AgNPs exhibit a strong absorption band and generate specific color in solution due to the surface plasmon resonance (SPR)^39^. At first, the colorless AgNO_3_ solution became light brown after a few hours, while it turned dark brown at the end of the synthesis. The brown color observed at the end of the synthesis might occur at the 400–500 nm wavelength range due to the stimulation of surface plasmon vibrations specific to AgNPs^40^.

The reduction of Ag^+^ ion to Ag^0^ during the reaction with the ingredients in CLE extract was observed by UV-Vis spectroscopy. The absorption peak for the CLE sample was found at 405 nm (Fig. 1a), which confirmed the formation of the desired AgNPs.

**Figure 1.**
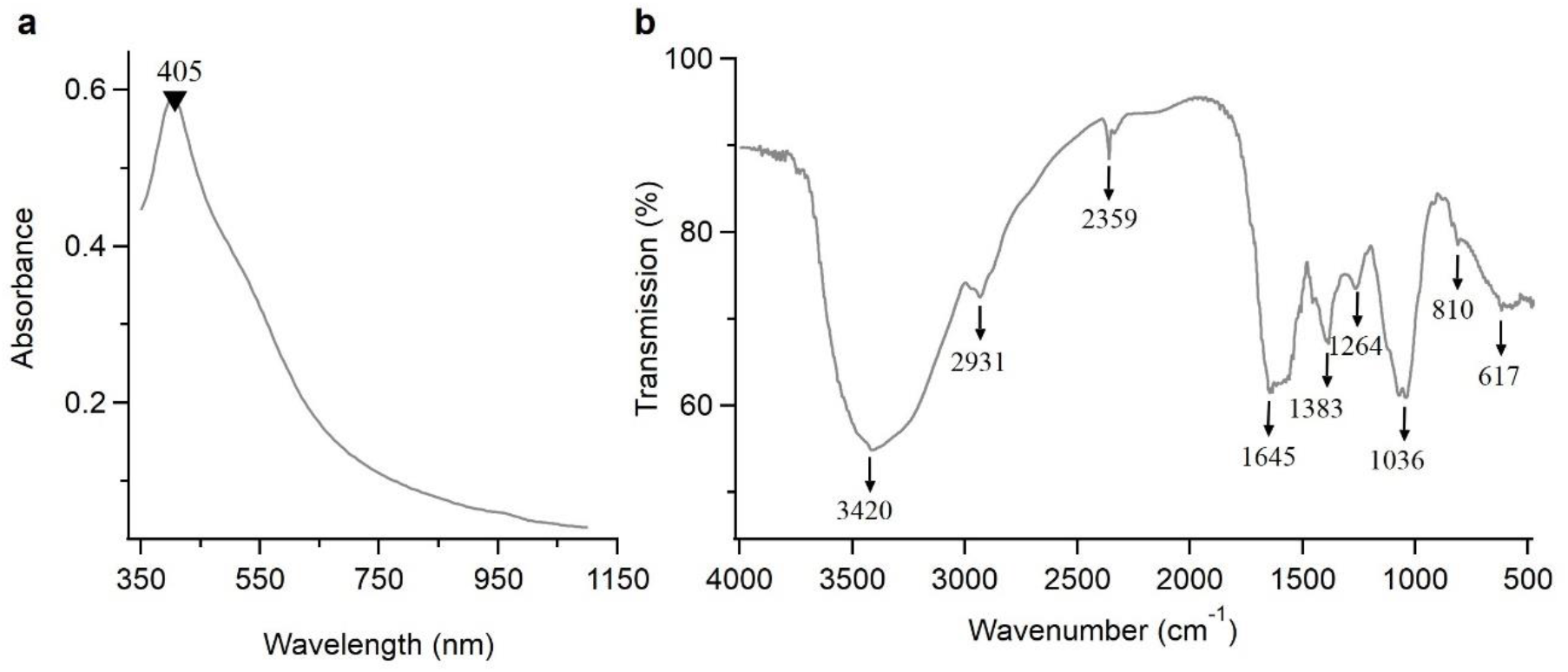
(a) UV-Visible spectra of AgNPs synthesized from *C. infortunatum* flower extract. (b) FTIR spectrum of synthesized AgNPs from aqueous extract of *C. infortunatum*.

#### Fourier transform infrared spectroscopy (FTIR) Analysis

To identify probable functional groups of the biomolecules present in Clerodendrum aqueous flower extract and responsible for the formation and stabilization of AgNPs, FTIR spectra of CLE AgNPs were recorded (Fig. 1b). The strong bands at 3420, 2931, 2359, 1645, 1383, 1264, 1036, 810 and 617 cm^−1^ corroborate with the capping agents responsible for the AgNPs formation. The vibration band at 3420 cm^−1^ in the spectra is assigned to O–H stretching in the molecules. The peaks near 2391 and 2359 cm^−1^ are assigned to C–H stretching and aldehydic O–H stretching, respectively. The weak band at 1645 cm^−1^ corresponds to C=C stretching arising due to the presence of carboxylic acids. The peak at 1383 cm^−1^ corresponds to the alkane’s C–O stretching vibration. The peak near 1264 cm^−1^ corresponds to C-stretching (nonconjugated). The peaks near 1036 cm^−1^ and 810 cm^−1^ are assigned to C–N and C–H stretching, respectively. The spectra also showed a sharp peak at 617 cm^−1^ corresponding to the C–Br (alkyl halides) stretching vibration.

#### X-ray diffraction (XRD) analysis

To identify the crystalline phase and further confirm the AgNPs formation, the X-ray diffraction pattern of silver nanoparticles was recorded at UGC-DAE Consortium for Scientific Research, Kolkata (Bruker d8 Advance X-ray diffractometer, CuKα radiation (λ = 1.5406 Å), 40 kV-40mA, 2θ/θ scanning mode). Data was collected for the 2θ range of 20 to 80 degrees with a step of 0.0202 degree. The XRD patterns display five characteristic peaks at 38.188, 44.364, 64.53, and 77.485 Å (Fig. 2). These peaks correspond to the crystal planes (111), (200), (220), and (311), respectively and match with the powdered diffraction standard values of Miller indices (*hkl*) of silver. The diffraction angles corresponded to the face-centered cubic (FCC) structure of silver in AgNPs and agreed with the standard powder diffraction card of the Joint Committee on Powder Diffraction Standards (JCPDS) corresponding to silver (File No.: 04–0783). The average crystal size *D* of the AgNPs has been estimated from the diffractogram by using the Debye-Scherrer formula, *D* = 0.9*λ*/*β* Cos *θ*, where *λ* is the wavelength of the X-rays used for diffraction, θ the Bragg angle, and *β* the full width at half maximum (FWHM) of a diffraction band^41^. Based on the XRD spectrum of the CLE-AgNP, the average size of the nanoparticles was calculated to be 27.67 nm.

**Figure 2.**
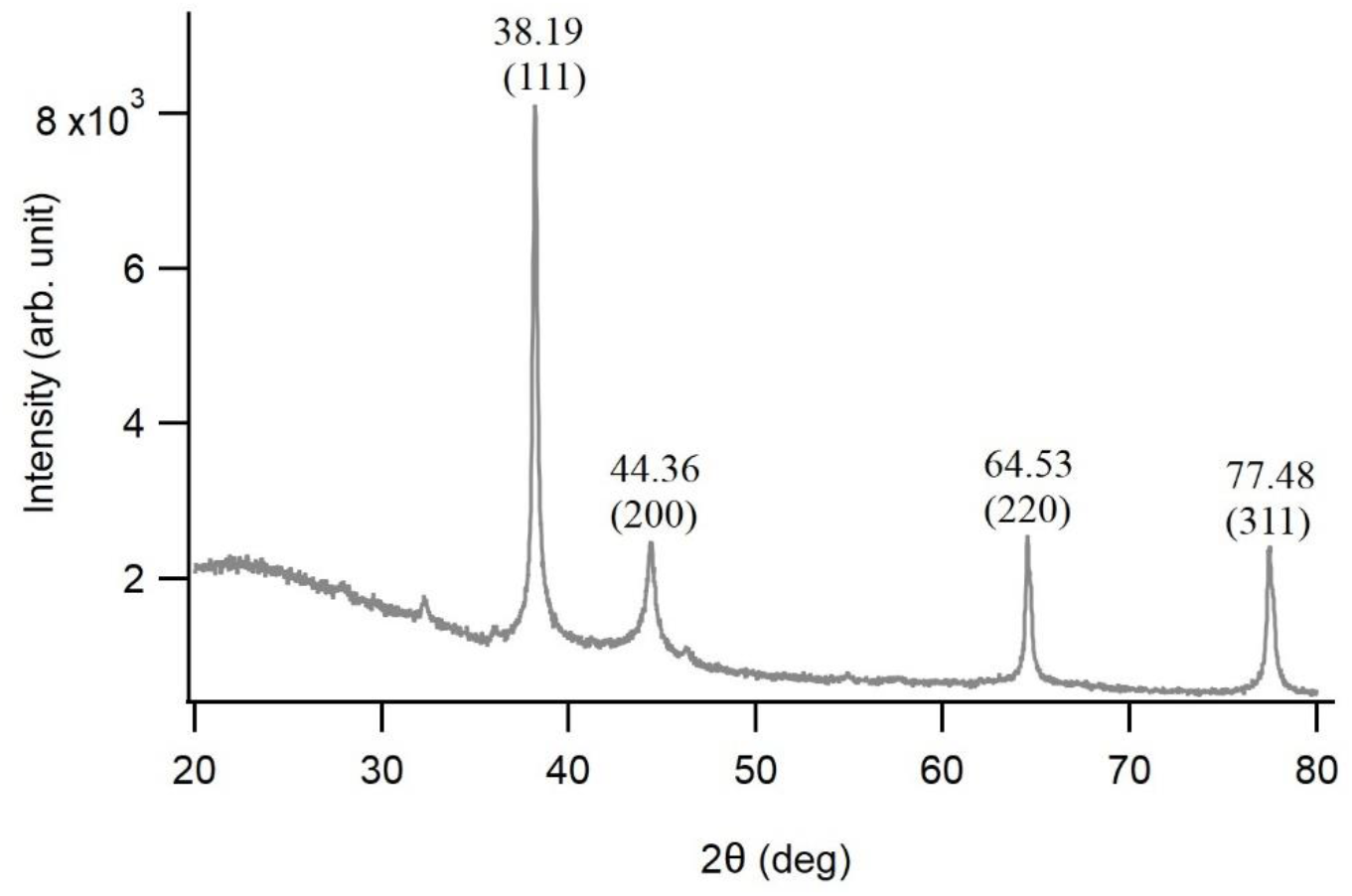
X-ray diffraction patterns of synthesized CLE-AgNPs.

#### Transmission Electron Microscopy (TEM) studies

The dispersion, aggregation, crystalline state and size of the CLE-AgNPs were examined using TEM, as shown in Fig. 3a. The sizes of AgNP particles were measured using ImageJ (https://imagej.nih.gov/ij/), and a histogram of sizes was calculated. The Gaussian fitting of the histogram yielded the mean particle size of AgNPs to be 27.75 ± 3.13 nm (mean ± sd, N = 90) (Fig. 3b). The surface morphology studied by TEM showed clear evidence for the metallic crystal formation of AgNPs, which was dispersed uniformly with fewer particle aggregation, and the particles formed in a spherical shape. The presence of some larger nanoparticles may be attributed to the fact that AgNPs tend to aggregate due to their high surface energy and high surface tension of the ultrafine nanoparticles^42^.

**Figure 3.**
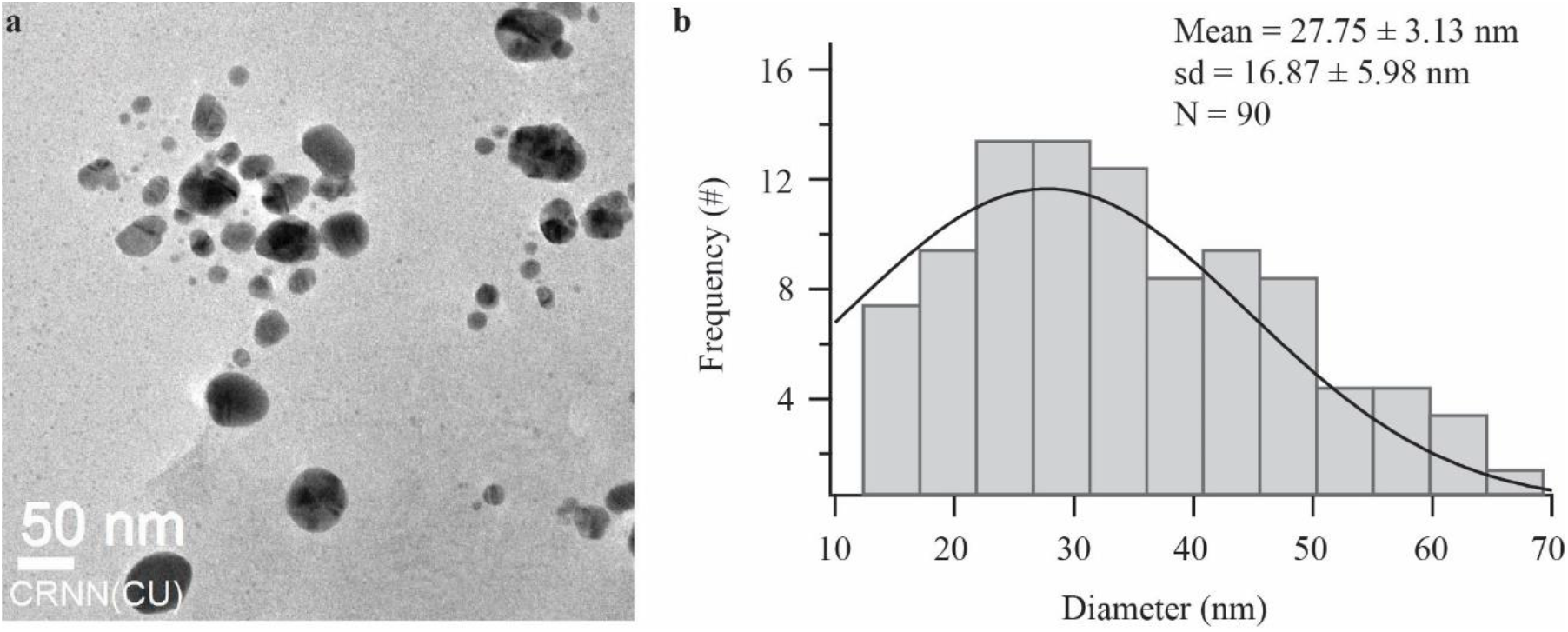
(a) A typical TEM micrographic image of synthesized AgNPs. (b) AgNPs size distribution extracted from TEM images. The solid black curve is a Gaussian fit to the data.

#### Efficacy of CLE-AgNP on *Raillietina* spp

*Raillietina* spp., incubated in the control medium (PBS only), showed physical activity for a longer period; the controls survived for about 72.00 ± 0.04 h, following which they became immobilized and dead (Fig. 4). On exposure to the test medium (CLE-AgNP and Genistein), the parasites proceeded from the vigorous movement state to the relaxed state, following which they attained paralysis leading to death. The *in vitro* evaluation of the efficacy of CLE-AgNPs against the cestode parasite showed a paralysis time within 1.51 h, 1.17 h, 0.59 h, 0.48 h, 0.43 h and death time of 2.48 h, 2.11 h, 1.41 h, 1.27 h, 1.07 h for dosages of 25, 50, 75, 100, and 125 µg/ml, respectively.

**Figure 4.**
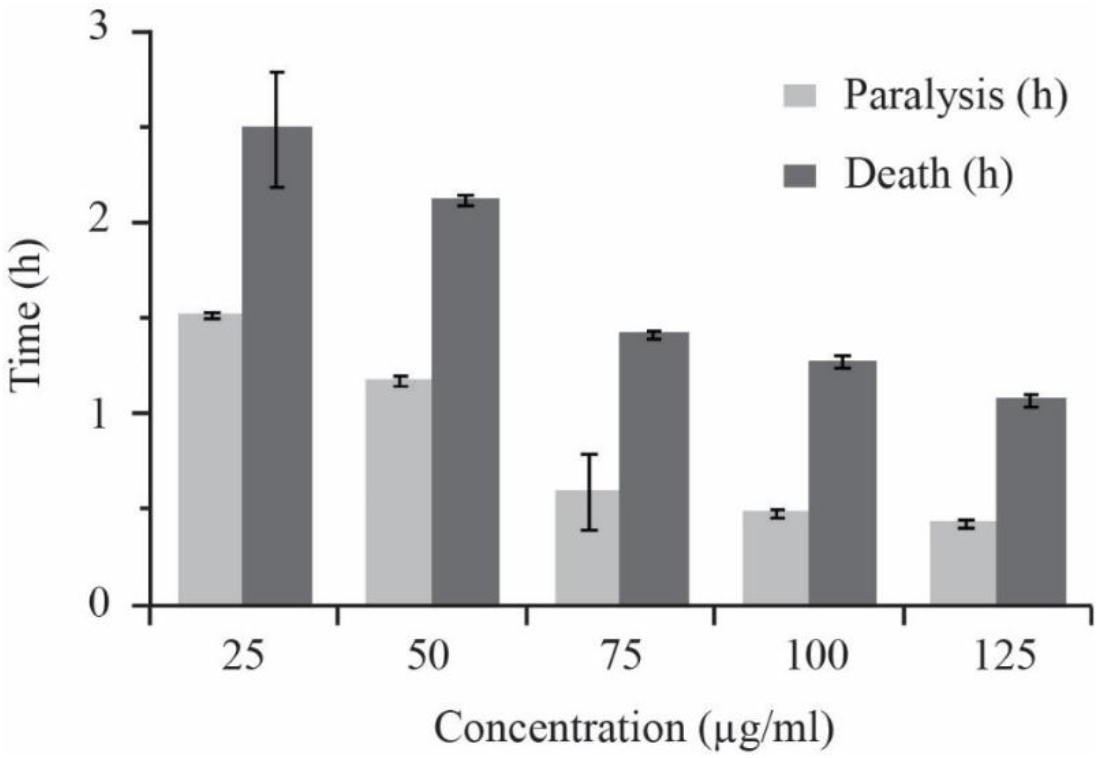
Results of CLE-AgNP efficacy on *Raillietina* spp. after exposure to five different concentrations (25, 50, 75, 100 and 125 µg/ml PBS).

#### Morphological changes of CLE-AgNP exposed *Raillietina* spp

SEM images of the control parasites reveal a rostellum and four suckers arranged sideways around the scolex, each with circlets of broad hooks at the bottom, tapered, and bent toward the ends (Fig. 5a). At higher magnification, the surface tegument of the proglottid is revealed to be covered with smooth, homogenous microtriches, which are the absorptive structures for feeding (Fig. 5d). However, on exposure to the test medium, the general surface topography of the proglottids degenerated with the formation of wrinkles (Fig. 5f). The cestode treated with CLE-AgNP showed irrevocable destruction of the scolex which appeared greatly distorted with suckers extensively shrunken and sharply crooked spines around the suckers. The filamentous nature of microtriches was lost, and the spines around suckers were broken and eroded (Fig. 5c), which altered the maintenance of the parasite position on the host cell, affecting its nutrition. The Genistein (reference drug) treated parasites showed immense disfigurement of the scolex, breakage and disembarkment of the tegumental surface structures (Figs. 5b, e).

**Figure 5.**
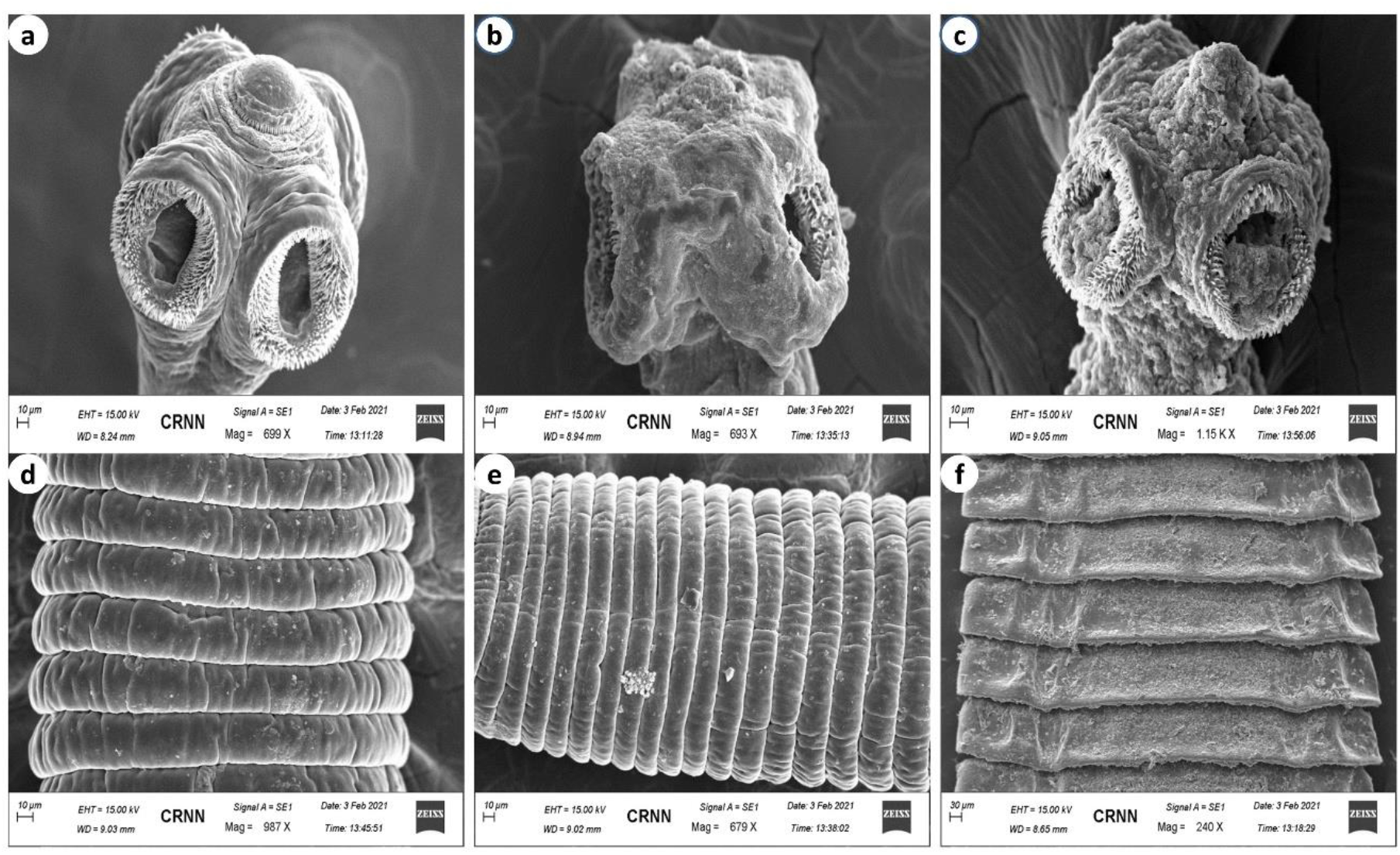
Scanning electron micrographs of control worm (a - scolex, d - gravid proglottid); Genistein (b - scolex, e - gravid proglottid) and silver nanoparticle exposed *Raillietina* spp. (c - scolex, f - gravid proglottid).

#### Histochemical studies

The tegument of the control *Raillietina* spp. showed intense activity of AcPase, AlkPase, ATPase and 5’-Nu compared to sub-tegument and somatic musculature (Figs. 6a-d). Figures 6e-h depict the histological sections of CLE-AgNP exposed parasites showing a general reduction in the staining intensity in the tegument, sub tegument and somatic musculature. AcPase showed a pronounced reduction in stain intensity in the tegument and sub-tegument region for the AgNP incubated section (Figs. 6a, e). However, there was minimal activity throughout the section of parasites exposed to Genistein (Figs. 6i-l). A much-diminished activity of AlkPase was observed throughout the treated sections of the parasite (Figs. 6b, f). The ATPase activity was almost imperceptible in the tegument and sub-tegument of the parasite treated with CLE-AgNP compared to the control parasites (Figs. 6c, g). The 5’-Nu activity was also found to be reduced throughout the tegument and sub-tegumental region in the AgNP exposed cestodes compared to the control (Figs. 6d, h).

**Figure 6.**
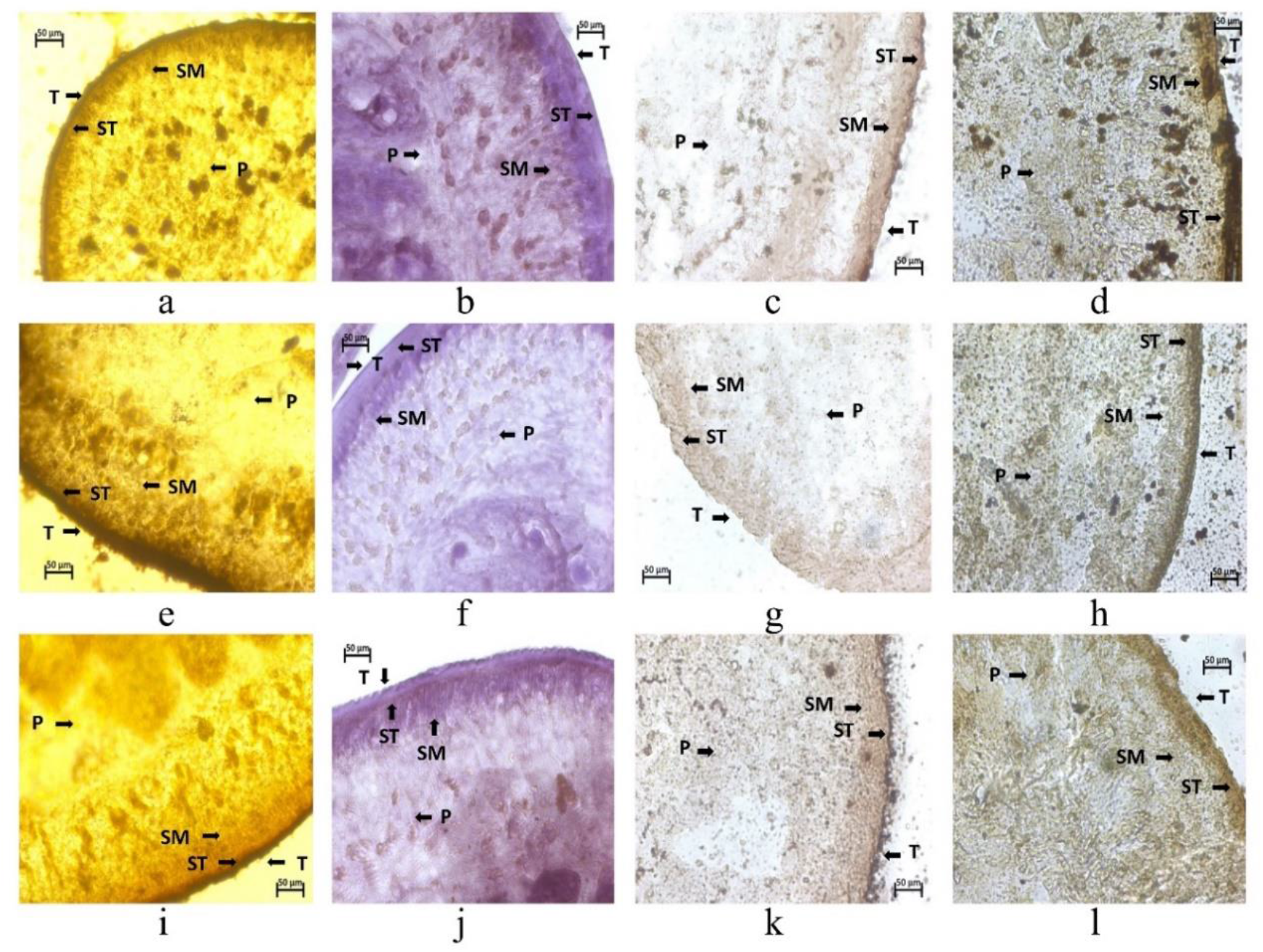
Histochemical demonstration of AcPase (a, e, i), AlkPase (b, f, j), ATPase (c, g, k) and 5’-Nu (d, h, l) activities in *Raillietina* spp. treated with (125 μg/ml) and Genistein GEN (125 μg/ml); (a-d): Transverse section of control parasite; (e-h): AgNPs-exposed parasite; (i-l): GEN-exposed parasite. All scale bars correspond to 50 μm.

## Discussion

In this study, we utilized plants for the extracellular production of silver nanoparticles and demonstrated their capabilities as an alternative to synthetic chemical techniques. In recent times, the synthesis of nanoparticles from natural sources has gained immense publicity. Organic nanoparticles such as chitosan and lipid nanoparticles and inorganic nanoparticles such as gold have been used as nano-drug delivery systems^43-47^. The mode of bactericidal effect of nanoparticles has not yet been fully elucidated. However, it irreversibly disrupts the bacterial cell wall, inactivates vital proteins, chelates DNA, and forms reactive oxygen species (ROS), known to have high microbicidal activity^47-51^. While microorganisms continue to be investigated for metal nanoparticle synthesis, using plant extracts in the biosynthesis of nanoparticle manufacturing processes is an attractive prospect that is yet largely unknown and underutilized. AgNPs play a profound role in biology and medicine due to their attractive physiochemical properties.

In the current investigation, we focused on the green synthesis of CLE-AgNP by adding plant extract to the AgNO_3_ solution, which was initially confirmed by the color changes of the solution (dark yellowish to brown) and later measured spectrophotometrically (Fig. 1a). In a similar study,

The FTIR measurements of CLE-AgNP showed broad absorption spectra for the hydroxyl group between 3309.85 cm^−1^, confirming the involvement of diterpenes (Fig. 1b). Further, the sharp peak at 1606.70 cm^−1^ indicated the presence of the phenyl group. The sharp peak at 1315.45 cm^−1^ indicated C(O)–O stretching vibrations; the peak at 1031.91 cm^−1^ indicated the involvement of reducing sugars^52^. Furthermore, Al Masoud *et al*. demonstrated the use of FTIR to detect biomolecules for reducing Ag+ ions and capping ginger extract-produced AgNPs^53^. The findings showed that the significant peaks at 1057, 1386, 1620, and 3417 cm−^1^ corresponded to primary alcohols and NO_3_ and C−N stretching vibrations of aliphatic and aromatic amines, respectively^54^. CLE-AgNP was crystalline, mostly spherical, with variable particle size and elemental (Figs. 2, 3a, b). The crystallite size of the material from XRD measurements was found to be around 27.67 nm). In addition to the four major XRD peaks in the diffractogram, other unidentified peaks observed might be attributed to the formation of the crystalline biomolecules bound to the surface of the AgNPs. TEM analysis also confirmed the synthesis of variable size (12–44 nm, mean ≈ 27) and spherical shape AgNPs (Figs. 3a, b). The results demonstrate that CLE-AgNP indeed has anthelmintic efficacy on poultry cestode *Raillietina* spp. and has acted in a dose and time-dependent manner. The most efficacious dose was 125 µg/ml, wherein the initiation of paralysis of parasites occurred after 0.43 h and death after 1.07 h of treatment (Fig. 4 and Supplementary Table S1). Scanning electron microscopy results have shown discernible topographical mutilation on test worms, while control tapeworm shows smooth and organized rostellum, sucker, and microtriches on the head and strobila portion. As observed in the present investigation, tegumental erosion has altered the host-parasite interface, causing severe nutrient deficiency within the parasite. Apart from the absorption of nutrients, the possible role of spine-like features of the microtriches is holding with its host to maintain its position in the gut. Therefore, disruption of microtriches leads to reduced attachment capacity of the parasites to the host. Similar observations were recorded on incubation of the parasite in Resveratrol, which causes blebbing, swelling of the tegument, loss of spines and distortion of suckers in *Raillietina* spp. exposed *in vitro*^28^. Extensive blebbing of *Fasciola hepatica* surface was seen on treatment with various concentrations of crude ethanolic shoot extract of traditionally used medicinal plants like *Alpinia nigra*^55^. Other phytocompounds like α-viniferin have been unequivocally responsible for damages in varying degrees on the teguments of cestodes^56^. AcPase and AlkPase are found to be involved in the uptake of certain nutrients, glycogen and lipoprotein in various helminth parasites^57^. Enzymatic alterations in histochemical localization studies also revealed pronounced effects (Table 1). A significant reduction in activity in the somatic musculature and complete absence in the tegument and sub-tegumental layers in 125 µg/ml AgNP exposed parasite further indicate the disruption of the energy metabolism pathway in the parasite. Following the treatment of Genistein, the active component of *Flemingia vestita*, the tegumental enzymes like AcPase, AlkPase, ATPase and 5’-Nu of *Raillietina* spp. were found to be decreased many folds^58^. Thus, the observed decrease in the two tegumental phosphatase activities might be related to inhibition or decreased glucose absorption by *Raillietina* spp., resulting in progressive loss of motor function owing to a deficiency of energy source and, eventually, paralysis and death.

**Table 1.**
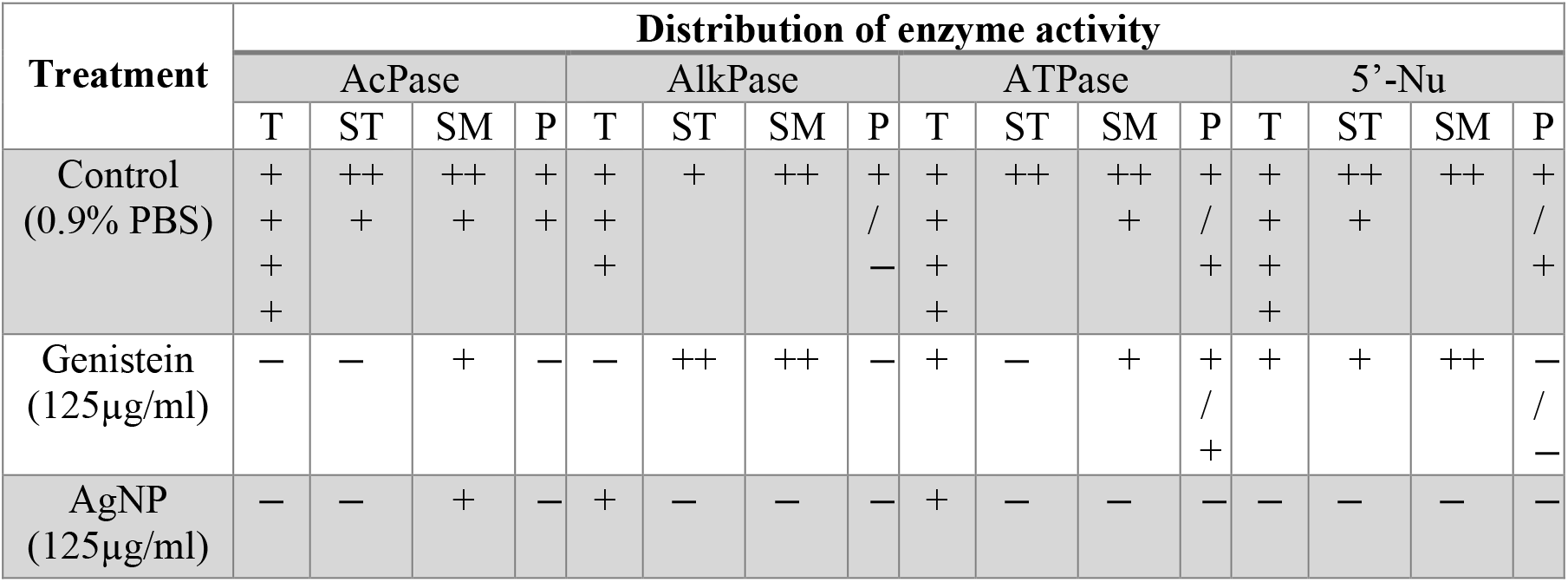
Summary of histochemical localization activities of AcPase, AlkPase, ATPase and 5’-Nu in the various structures of *Raillietina* spp. T = Tegument, ST = Sub-Tegument, SM = Somatic Musculature, P = Parenchyma. + + ++ = very intense activity, + + + = intense activity, + = mild activity, − = no activity.

Our investigation elucidates a concise, cost-effective, and productive route in the phytosynthesis of AgNPs utilizing the aqueous floral extract of CLE. Further in vivo studies are required to understand the mode of action of the drug and its pharmacokinetics. In the future, optimization of the test therapeutic (CLE-AgNP) through pharmaceutical evaluation and application of appropriate feeding strategy by finding out the effects on metabolic precursors and intracellular metabolites can contribute to a seamless transition to financially feasible ethnomedicinal anthelmintic. We believe using green synthesis methods to produce silver nanoparticles from *Clerodendrum infortunatum* flowers could provide a promising alternative to synthetic anthelmintic drugs for treating helminth infections in domestic fowl.

## Materials and Methods

### Collection of plant material

The fresh and healthy flowers of *Clerodendrum infortunatum* were collected from the Cooch Behar Panchanan Barma University (CBPBU) campus garden (latitude 26.321796 °N, longitude 89.469329 °E). Dr. Chaya Deori did the taxonomical evaluation, and the voucher specimen (AC-97296) was kept at the herbarium of the Botanical Survey of India, Eastern Regional Centre, Shillong.

### Preparation of plant extract using flowers of CLE

The flowers were thoroughly washed sequentially with tap and deionized water, air-dried and chopped into small pieces. Chopped flowers (20 g) were added to 100 ml deionized autoclaved water in a beaker and heated at 90°C in a temperature-controlled water bath for 10 min. The extract was cooled, filtered through Whatman No. 1 filter paper, and kept at 4°C until used.

### Biosynthesis of CLE-capped AgNP

For silver nanoparticle synthesis, about 10 mL of CLE flower extract was added to a 90 mL aqueous solution of 1 mM AgNO_3_ (Merck Laboratories, India) and kept at room temperature. The color changed from pale yellow to brown, indicating the formation of the AgNPs due to the reaction of flower extracts of CLE with silver metal ions. Control was maintained without adding flower extract in the silver nitrate solution that showed no color changes. The purified AgNPs were obtained by removing the extract by centrifuging the suspension thrice at 15,000 rpm for 20 min, followed by washing it twice with double sterilized water.

### Characterization studies of AgNP

The bio-reduction process of silver ions in solution was preliminarily determined by visible colour changes and later monitored using UV–visible absorption spectroscopy. The synthesized AgNPs were freeze-dried, powdered, and used for X-ray diffraction (XRD) analysis. The dried powder was mixed with potassium bromide at the ratio of 1:100, and the results were recorded using the Fourier transform infrared spectroscopy (FTIR). Transmission electron microscopy (TEM) was used to understand the size and morphology of AgNPs.

### Collection of parasites and *in vitro* treatment

Live mature *Raillietina* spp. (Megnin, 1880) (Class: Cestoda; Order: Cyclophyllidea; Family: Davaineidae) were collected from the intestine of freshly sacrificed domestic fowl (*Gallus gallus domesticus* L.) from local abattoirs in Cooch Behar and maintained in 0.9% PBS at 37 ± 1C in an incubator. Control parasites were maintained in 0.9% PBS (pH 7.2) at 37 ± 1°C, whereas for treatment, live worms were directly incubated in different concentrations of test treatment (25, 50, 75, 100, 125 µg/ml) in separate Petri dishes. Similarly, treatment was also performed with Genistein (GEN) at a dose of 125 µg/ml of PBS as a broad-spectrum reference drug. Six replicates for each set of incubation mediums were prepared, and the time taken to attain the paralytic state and death was recorded. Mortality of parasites was confirmed by removing treated parasites from the test medium and dipping them in slightly lukewarm water, which was indicated by the cessation of all signs of movement. The treated and control parasites were retrieved from the respective incubation mediums and processed for histochemical and electron microscopic studies.

### Scanning Electron microscopy (SEM) of the parasite

The parasites were fixed in 10% neutral buffered formalin (NBF) for 24 hours immediately after paralysis. After fixation, the sample was rinsed in PBS and dehydrated with acetone grades ranging from pure dry acetone to escalating degrees of acetone. The specimens were then critical-point dried using liquid carbon dioxide as the transitional fluid, which was put on a metal stub and coated with platinum in a fine-coat ion sputter, JFC-1100 (JEOL). The specimens were then viewed using a Zeiss EVO-18 (Special edition) scanning electron microscope at an accelerating voltage of 10–15 kV.

### Histochemical studies

The tegumental enzymes were investigated histochemically using duly processed frozen sections cut at a thickness of 10–12 µm in a Leica CM 3050S cryostat. Acid phosphatase (AcPase) activity was detected in cold formol calcium fixed specimens following the modified lead nitrate method^59^, using sodium β-phosphoglycerate as the substrate, where a brownish precipitate on tegumental sections indicates the sites of AcPase activity. The modified coupling azo-dye method was used to assess alkaline phosphatase (AlkPase) activity at room temperature (17–20 °C). The calcium method of Pearse was implemented to detect the localization of adenosine triphosphatase (ATPase) activity, where adenosine triphosphate (ATP) was used as the substrate, and the enzyme activity was determined by observing blackish-brown deposits^59^. To observe 5’-Nucleotidase (5’-Nu), the lead method of Wachstein and Meisel was employed using adenosine monophosphate as a substrate^60^.

## Acknowledgment

The authors express their heartfelt gratitude to Dr. Debkumar Mukhopadhyay, Vice-Chancellor of CBPBU, for his support and encouragement and for providing the necessary laboratory facilities. The authors are also grateful to the UGC-DAE Consortium, Kolkata, for the XRD facility and Centre of Nanoscience and Nanotechnology (CRNN), Kolkata, for allowing us to use their SEM, TEM and FTIR facility.

## Author contributions

RM and PKK conceived the idea and designed the research. RM performed the experiments, analyzed, and interpreted the data. RM and PKK wrote the manuscript.

## Availability of data and material

The data used to substantiate the findings of this study are included in the article; however, the raw data is also available from the corresponding author upon request.

## Additional Information

### Ethics approval

We confirm that any aspect of the work covered in this manuscript does not involve ethical approval.

### Consent for publication

We affirm that all authors read and appr’oved the manuscript and that no other individuals satisfied the authorship requirements but are not included. We further confirm that all have approved the order of the authors listed in the manuscript of us.

