## Supplemental Table 1 for "Biosynthesis, Characterization and Anthelmintic Activity of Silver Nanoparticles of *Clerodendrum infortunatum* Isolate"

| Test material | Concentration (µg/ml PBS) | Paralysis (h) | Death (h) |
|---|---|---|---|
| **Control** | _ | _ | 72 ± 0.04 |
| **CLE-AgNP** | 25 | 1.51 ± 0.02 | 2.48 ± 0.30 |
| | 50 | 1.17 ± 0.03 | 2.11 ± 0.03 |
| | 75 | 0.55 ± 0.20 | 1.41 ± 0.02 |
| | 100 | 0.48 ± 0.02 | 1.27 ± 0.03 |
| | 125 | 0.43 ± 0.02 | 1.07 ± 0.03 |
| **Genistein** | 125 | 0.49 ± 0.02 | 1.33 ± 0.02 |

Supplementary Table S1. In addition to the *in vitro* efficacy of CLE-AgNPs as shown in Fig. 4, the *Raillietina* spp. parasites kept in control and reference drug Genistein showed dose-dependent anthelmintic activity at all concentrations tested.
